# The Vitamin D Receptor (VDR) Regulates Mitochondrial Function in C2C12 Myoblasts

**DOI:** 10.1101/872127

**Authors:** Stephen P. Ashcroft, Joseph J. Bass, Abid A. Kazi, Philip J. Atherton, Andrew Philp

## Abstract

Vitamin D deficiency has been linked to a reduction in skeletal muscle function and oxidative capacity, however, the mechanistic basis of these impairments are poorly understood. The biological actions of vitamin D are carried out via the binding of 1α,25-dihydroxyvitamin D3 (1α,25(OH)_2_D_3_) to the vitamin D receptor (VDR). Recent evidence has linked 1α,25(OH)_2_D_3_ to the regulation of skeletal muscle mitochondrial function *in vitro*, however, little is known with regard to the role of the VDR in this process. To examine the regulatory role of the VDR in skeletal muscle mitochondrial function, we utilised lentiviral mediated shRNA silencing of the VDR in C2C12 myoblasts (VDR-KD) and examined mitochondrial respiration and protein content compared to shRNA scrambled control. VDR protein content was reduced by ~95% in myoblasts and myotubes (*P* < 0.001). VDR-KD myoblasts displayed a 30%, 30% and 36% reduction in basal, coupled and maximal respiration respectively (*P* < 0.05). This phenotype was maintained in VDR-KD myotubes, displaying a 34%, 33% and 48% reduction in basal, coupled and maximal respiration (*P* < 0.05). Furthermore, ATP production derived from oxidative phosphorylation (ATP_ox_) was reduced by 20% suggesting intrinsic impairments within the mitochondria following VDR-KD. However, despite the observed functional decrements, mitochondrial protein content as well as markers of fusion and fission were unchanged. In summary, we highlight a direct role for the VDR in regulating skeletal muscle mitochondrial respiration *in vitro*, providing a potential mechanism as to how vitamin D deficiency might impact upon skeletal muscle oxidative capacity.

## INTRODUCTION

Vitamin D deficiency is characterised by serum 25-hydroxyvitamin D (25(OH)D) levels of <50 nmol.L^−1^ (14). Based upon these numbers, it has been reported that approximately 40% of adults in the USA can be classified as deficient (6). The classical actions of vitamin D are well established, primarily functioning to maintain calcium and phosphate balance in order to prevent bone related disease (1, 12). Vitamin D carries out its actions via its active metabolite, 1α,25-dihydroxyvitamin D3 (1α,25(OH)_2_D_3_), which binds to the ubiquitously expressed vitamin D receptor (VDR) (13). The VDR, together with its binding partner retinoid x receptor alpha (RXRα), recruit transcriptional cofactors to regulate genomic transcription (16, 19).

In addition to its role in bone biology, vitamin D has also been shown to play a role in skeletal muscle development (10, 18) and regeneration (18). Given that vitamin D exerts its biological actions through binding to the VDR, multiple studies have sought to elucidate the role of the VDR within skeletal muscle (8, 9, 11). For example, whole body VDR knockout mice (VDRKO) present muscle weakness, muscle fibre atrophy and hyper-nuclearity (8), which is also present in muscle-specific VDR knock-out (VDR-mKO) mice (9). Collectively these studies suggest a specific role for the VDR in skeletal muscle regulation (3, 9).

In addition to regulating skeletal muscle mass and function, evidence also suggests that vitamin D may regulate skeletal muscle mitochondrial function (2, 24). For example, treating human primary myoblasts with 1α,25(OH)_2_D_3_ resulted in an improvement in mitochondrial function and an increase in ~80 mRNAs encoding for mitochondrial proteins (21). In addition, the VDR appeared to be critical in mediating the effects of 1α,25(OH)_2_D_3_, as siRNA targeted towards the VDR blocked mitochondrial adaptation. Therefore, the aim of the present work was to further examine the regulatory role of the VDR for mitochondrial function in skeletal muscle. To achieve this, we generated a stable VDR loss-of-function C2C12 cell line model and examined mitochondrial respiration and protein content in myoblasts and fully differentiated myotubes.

## METHODS

### Generation of VDR-KD and control cell lines

The lentiviral plasmid used (pLKO.1 backbone) was designed in-house and was based on (Clone ID: RMM3981-201757375) and targeted the (3’ UTR) mouse sequence 5′-TTA AAT GTG ATT GAT CTC AGG-3′ of the mouse *Vdr* gene; the scramble shRNA was used as a negative control as previously reported (15) with a hairpin sequence: CCT AAG GTT AAG TCG CCC TCG CTC TAG CGA GGG CGA CTT AAC CTT AGG (Addgene plasmid 1864, Cambridge, MA, USA). Oligos were obtained from ITDDNA USA (Integrated DNA Technologies, Inc. Iowa, USA) and suspended, annealed and cloned into pLKO.1 at EcoRI and AgeI restriction sites as per the pLKO.1 protocol from Addgene. The resultant plasmids were transformed in DH5α cells for amplification and isolated. The actual DNA sequence was confirmed at the Pennsylvania State University College of Medicine DNA sequence core facility. Packaging plasmids psPAX2 and envelope protein plasmid pMD2.G were a gift from Prof. Didier Trono, available as Addgene plasmids 12260 and 12259 respectively. HEK293FT cells (Invitrogen, Carlsbad, CA, USA) were grown in DMEM; 80–85% confluent plates were rinsed once with Opti-MEM (Invitrogen, Carlsbad, CA, USA) and then incubated with Opti-MEM for 4 h before transfections. psPAX2 and pMD2.G along with either scramble or pLKO.1 clones targeting mouse Vdr. Three clones were added after mixing with Lipofectamine 2000 as per the manufacturer’s instructions (Invitrogen, Carlsbad, CA, USA). Opti-MEM was changed after overnight incubation with DMEM containing 10% fetal bovine serum (FBS) without antibiotics to allow cells to take up the plasmids and recover. Culture media were collected at 36 and 72 h post-transfection for viral particles. Viral particles present in the supernatant were harvested after a 15-minute spin at 1,500 *g* to remove cellular debris. The supernatant was further filtered using a 0.45-μm syringe filter. Supernatant-containing virus was either stored at −80°C for long-term storage or at 4°C for immediate use. C2C12 myoblasts (ATCC, Virginia, USA) at 60% confluence were infected twice overnight with 3 ml of viral supernatant containing 8 μg.ml^−1^ polybrene in serum-free–antibiotic-free DMEM. Fresh DMEM media containing 10% FBS, 1% penicillin-streptomycin and 2 μg.ml^−1^ puromycin dihydrochloride (Sigma, St. Louis, MO, USA) were added the next day. Cells that survived under puromycin selection were harvested as stable VDR knock-down (VDR-KD) myoblasts or controls and stored in liquid N_2_ until further analysis.

### Extracellular flux analysis

Both control and VDR-KD cells were seeded in XFe24-well cell culture microplates (Seahorse Bioscience, North Billerica, MA, USA) at 3.0 × 10^5^ cells/well in 100 μl of growth medium. For myoblast experiments, cells were incubated at 37°C and 5% CO_2_ for 3 h in order to allow sufficient time for adherence and subsequently assayed. For myotube experiments, cells were incubated for a period of 24 h and medium changed to differentiation medium (DMEM, 2% horse serum and 1% pencilin-streptomycin). Differentiation media was changed every other day for 7 days. Prior to the assay, cells were washed and placed in 500 μl of Seahorse XF Base Medium (glucose 10 mM, sodium pyruvate 1 mM, glutamine 1 mM, pH 7.4) pre warmed to 37°C. The plate was then transferred to a non-CO_2_ incubator for 1 h. Following calibration, cell respiratory control and associated extracellular acidification were assessed following the sequential addition of oligomycin (1 μM), carbonyl cyanide *p*-trifluoromethoxyphenylhydrazone (1 μM) and a combination of antimycin A and rotenone (1 μM). Upon completion of the assay, cells were collected in sucrose lysis buffer (50 mM Tris pH 7.5; 270 mM sucrose; 1 mM EDTA; 1 mM EGTA; 1% Triton X-100; 50 mM sodium fluoride; 5 mM sodium pyrophosphate decahydrate; 25 mM beta-glycerolphosphate; 1 cOmplete™ protease inhibitor cocktail EDTA free tablet) and protein concentrations determined using the DC protein assay (Bio-Rad, Hercules, CA). Oxygen Consumption Rate (OCR) is reported relative to protein content (pmol/min/μg). Estimations of ATP production derived from both oxidative phosphorylation and glycolysis were performed as previously described (17).

### Mitochondrial Membrane Potential

Control and VDR-KD cells were plated at 1.0 × 10^5^ cells/well in 100 μl of growth medium in a black 96-well plate with a clear bottom (Corning, Costar, NY, USA). Cells were subsequently incubated for 30 minutes with 100 nM of tetramethylrhodamine ethyl ester (TRME). Following incubation cells were washed with PBS/0.2% BSA and then read at 549 nm using a CLARIOstar microplate reader (BMG Labtech, Germany) in 100 μl of PBS/0.2% BSA.

### Immunoblotting

Control and VDR-KD cells were plated at 1.0 × 10^10^ cells/well in 2 ml of growth medium in 6-well plates (Nunc, Roskilde, Denmark). Both myoblasts and myotubes were maintained and harvested as described previously, with protein concentrations determined using the DC protein assay (Bio-Rad, Hercules, CA). Total protein lysates of a known concentration were mixed 3:1 with 4x Laemmli sample loading buffer. Prior to gel loading, samples were boiled for 5 minutes unless probing for MitoProfile OXPHOS antibody cocktail, in which case non-denatured samples were used. The immunoblotting procedure was performed as previously described (25).

### Antibodies

All primary antibodies were used at a concentration of 1:1000 in TBS-T. Antibody for dynamin-1-like protein (DRP1;8570) was from Cell Signaling Technology; MitoProfile OXPHOS antibody cocktail (110413) and mitofilin (110329) were from Abcam; Optic Atrophy-1/dynamin-like 120 kDa protein (OPA1; CPA3687) was from BD Biosciences; citrate synthase (CS; SAB2701077) and mitochondrial fission protein 1 (FIS1; HPA017430) were from Sigma Aldrich; vitamin D receptor (D-6) (VDR; 13133) was from Santa Cruz Biotechnology. Secondary antibodies were used at a concentration of 1:10,000 in TBS-T. Anti-mouse (7076) and anti-rabbit (7074) were from Cell Signaling Technology.

### Statistical Analysis

Statistical analysis was performed using the Statistical Package for the Social Sciences (SPSS) version 24.0. Differences between control and VDR-KD C2C12s were determined by independent t-tests. All data is presented as mean ± standard deviation (SD). Statistical significance was set at *P* < 0.05.

## RESULTS

### Successful generation of VDR-KD myoblasts

Following shRNA interference, VDR protein content was reduced by 96% (*P* < 0.001) and 95% (*P* < 0.001) in VDR-KD C2C12 myoblasts (Fig. 1A) and myotubes (Fig. 1B) respectively.

**Figure 1.**
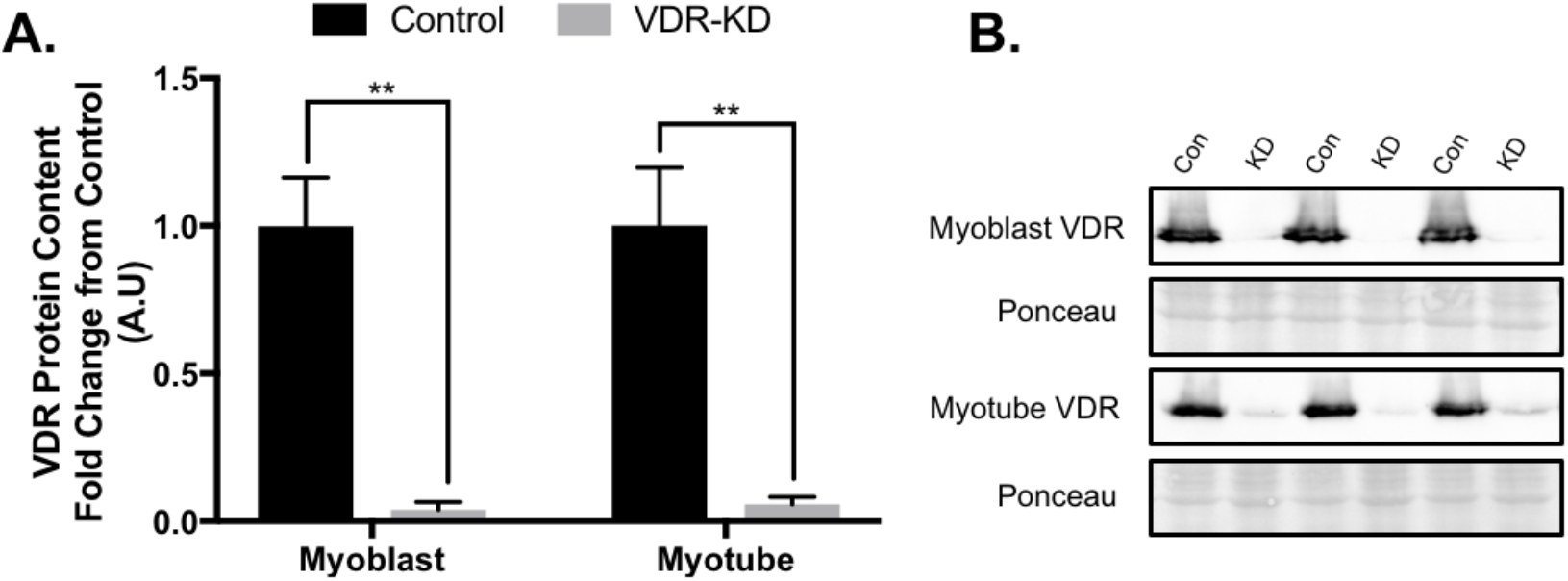
Generation of Vitamin D Receptor (VDR) loss of function C2C12 myoblasts. A: Quantification of VDR protein content in VDR-KD compared to control myoblasts and myotubes. B: Representative immunoblot images of VDR protein content in VDR-KD myoblasts and myotubes. ^**^*P* < 0.005, independent t-test. Data mean ± SD (n=5-6/group).

### VDR-KD results in reduced mitochondrial respiration in C2C12 myoblasts and myotubes

In order to determine the effects of VDR-KD upon mitochondrial function, extracellular flux analysis was performed in VDR-KD myoblasts and myotubes. VDR-KD myoblasts displayed a 30% reduction in basal respiration compared to control (*P* = 0.034; Fig. 2B). In addition, coupled and maximal respiration was reduced by 30% (*P* = 0.023) and 36% (*P* = 0.013) respectively, whilst spare respiratory capacity was also reduced by 39% (*P* = 0.008; Fig 2.B). This deficit was retained following differentiation, with VDR-KD myotubes displaying a 34% reduction in basal respiration (*P* < 0.001) and a 33% reduction in coupled respiration (*P* < 0.001) (Fig. 2D). Furthermore, maximal respiration was reduced by 48% (*P* < 0.001) and the spare respiratory capacity by 53% (*P* < 0.001; Fig. 2D) in VDR-KD. Whilst proton leak remained unchanged in VDR-KD myoblasts (Fig. 2B), VDR-KD myotubes displayed a 67% decrease in proton leak (*P* < 0.001; Fig. 2D). To establish where mitochondrial impairments originated, we estimated oxidative phosphorylation (ATP_ox_) and glycolysis (ATP_glyc_) using recently described equations (17). Accordingly, total ATP production and ATP_ox_ were reduced by 18% (*P* = 0.002) and 20% (*P* = 0.007) respectively in VDR-KD myoblasts (Fig. 2E). Finally, mitochondrial membrane potential assessed via TMRE fluorescence was reduced by 25% in VDR-KD (*P* = 0.001; Fig. 2F).

**Figure 2.**
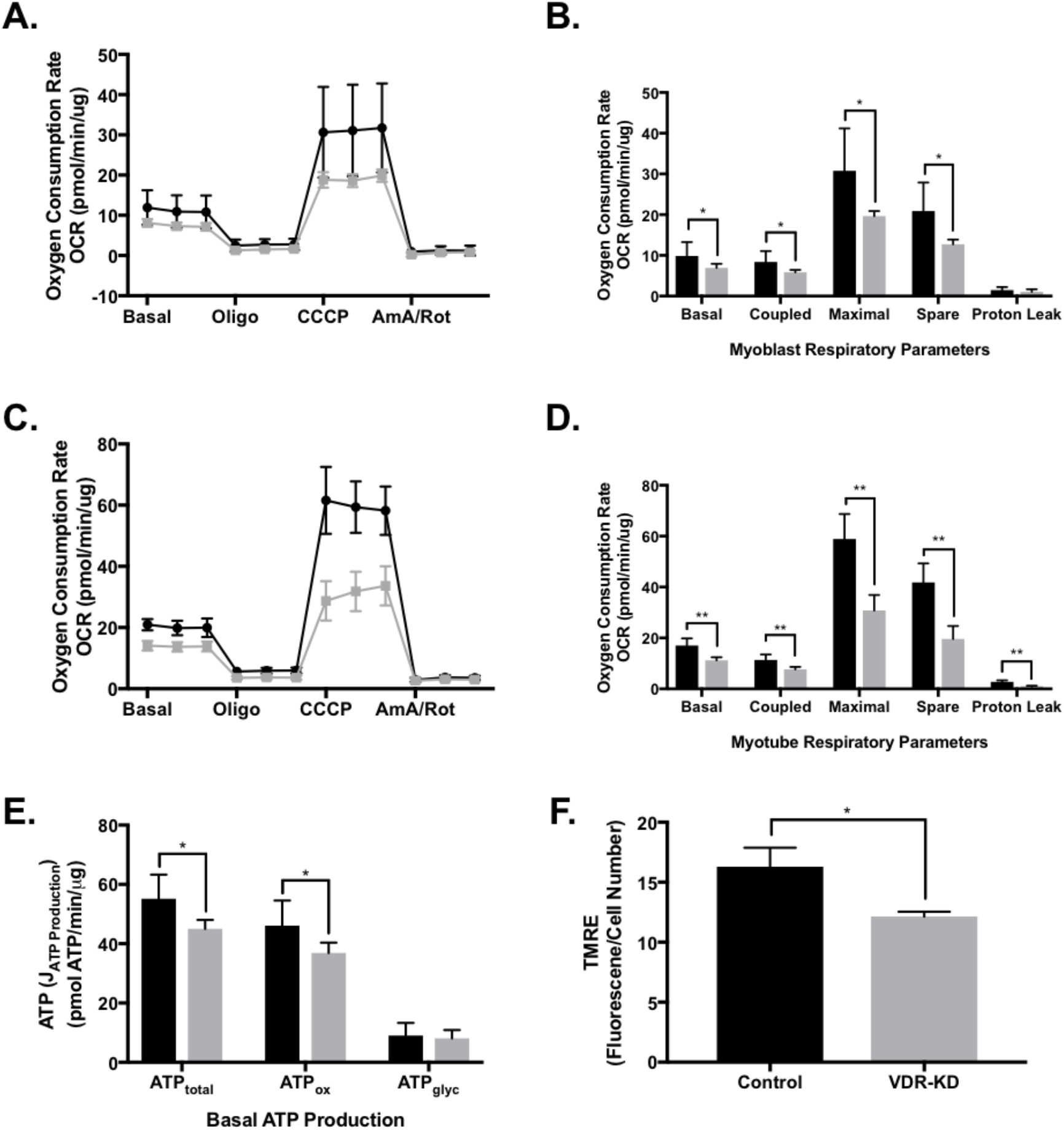
VDR-KD myoblasts display reduced mitochondrial respiration compared to control. A: Oxygen consumption rate (OCR) during analysis of respiratory control in control and VDR-KD myoblasts. B: Respiratory control parameters from control and VDR-KD myoblasts. C: OCR during analysis of respiratory control in control and VDR-KD myotubes. D: Respiratory control parameters from control and VDR-KD myotubes. E: Estimations of total ATP production (ATP_total_), oxidative phosphorylation (ATP_ox_) and glycolysis (ATP_glyc_) in control and VDR-KD myoblasts. F: Mitochondrial membrane potential assessed via TMRE fluorescence in control and VDR-KD myoblasts. ^**^*P* < 0.05, ^**^*P* < 0.005, independent t-test. Data mean ± SD (A-E: n=9-10/group. F: n=5/group).

### No change in mitochondrial related protein content in VDR-KD myoblasts and myotubes

Given the observed decrements in mitochondrial respiration in both VDR-KD myoblasts and myotubes, we sought to determine whether a reduction in mitochondrial related protein content might underlie this phenotype. However, no differences were observed in mitochondrial ETC subunit I-V, citrate synthase (CS) or cytochrome c (cyt c) protein content in either VDR-KD myoblasts (Fig. 3A) or myotubes (Fig. 3C). In order to further explore the potential influence of mitochondrial dynamics in mediating the observed decrements in mitochondrial function, multiple proxy markers of mitochondrial fusion and fission were probed. Whilst MFN2 remained unchanged, OPA1 increased by 15% in both VDR-KD myoblasts (*P* = 0.021; Fig. 4A) and myotubes (*P* = 0.046; Fig. 4C). Furthermore, Mitofilin, FIS1 and DRP1 all remained unchanged in VDR-KD myoblasts (Fig. 4A) and myotubes (Fig. 4C).

**Figure 3.**
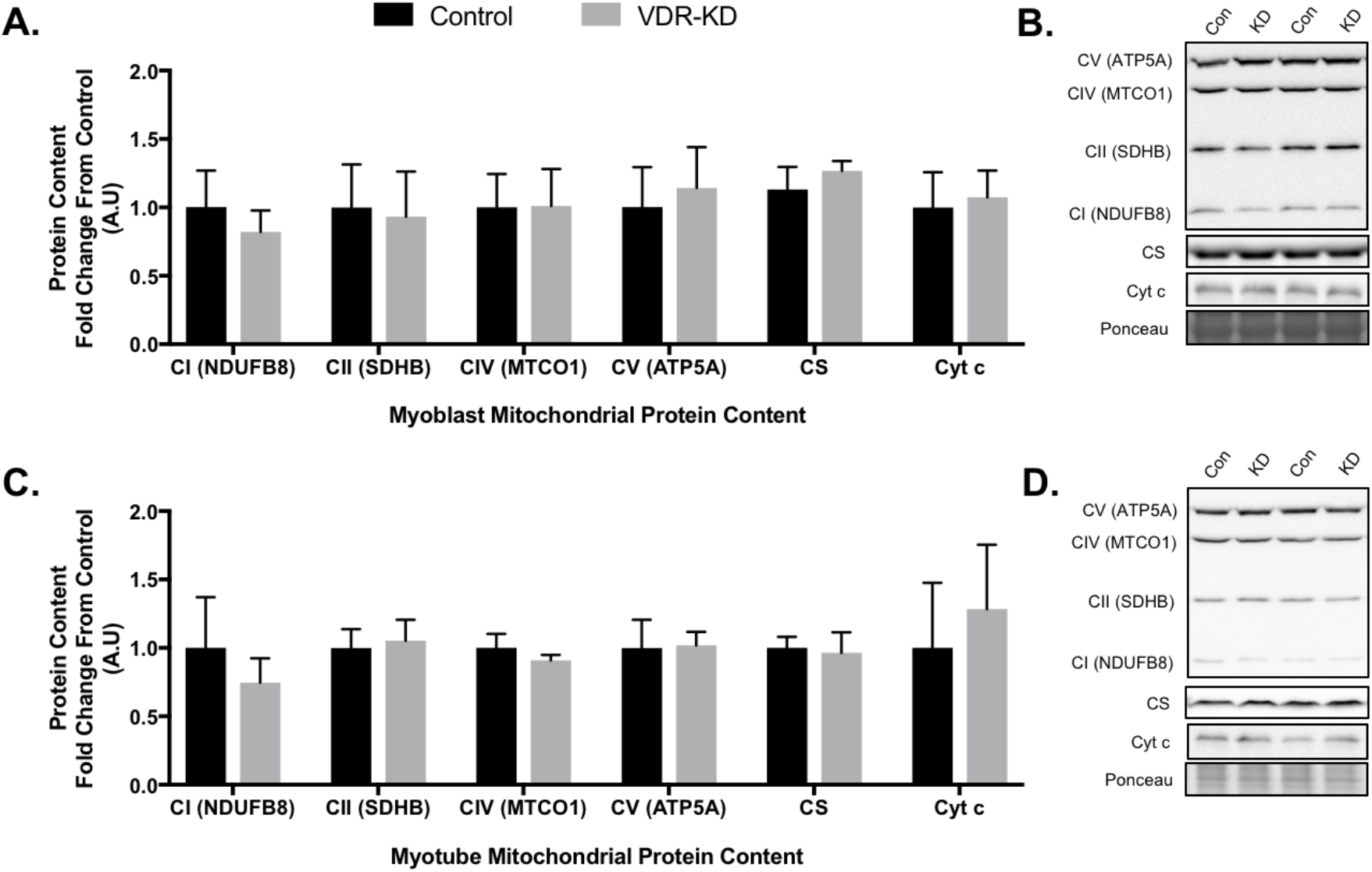
No change in markers of mitochondrial protein content in VDR-KD myoblasts compared to control. A: Protein abundance of mitochondrial subunits complex I (NDUFB8), complex II (SDHB), complex IV (MTCO1), complex V (ATP5A) as well as citrate synthase (CS) and cytochrome c (cyt c) in control and VDR-KD myoblasts. Data mean ± SD (n=6/group) and represented as a fold change from control.

**Figure 4.**
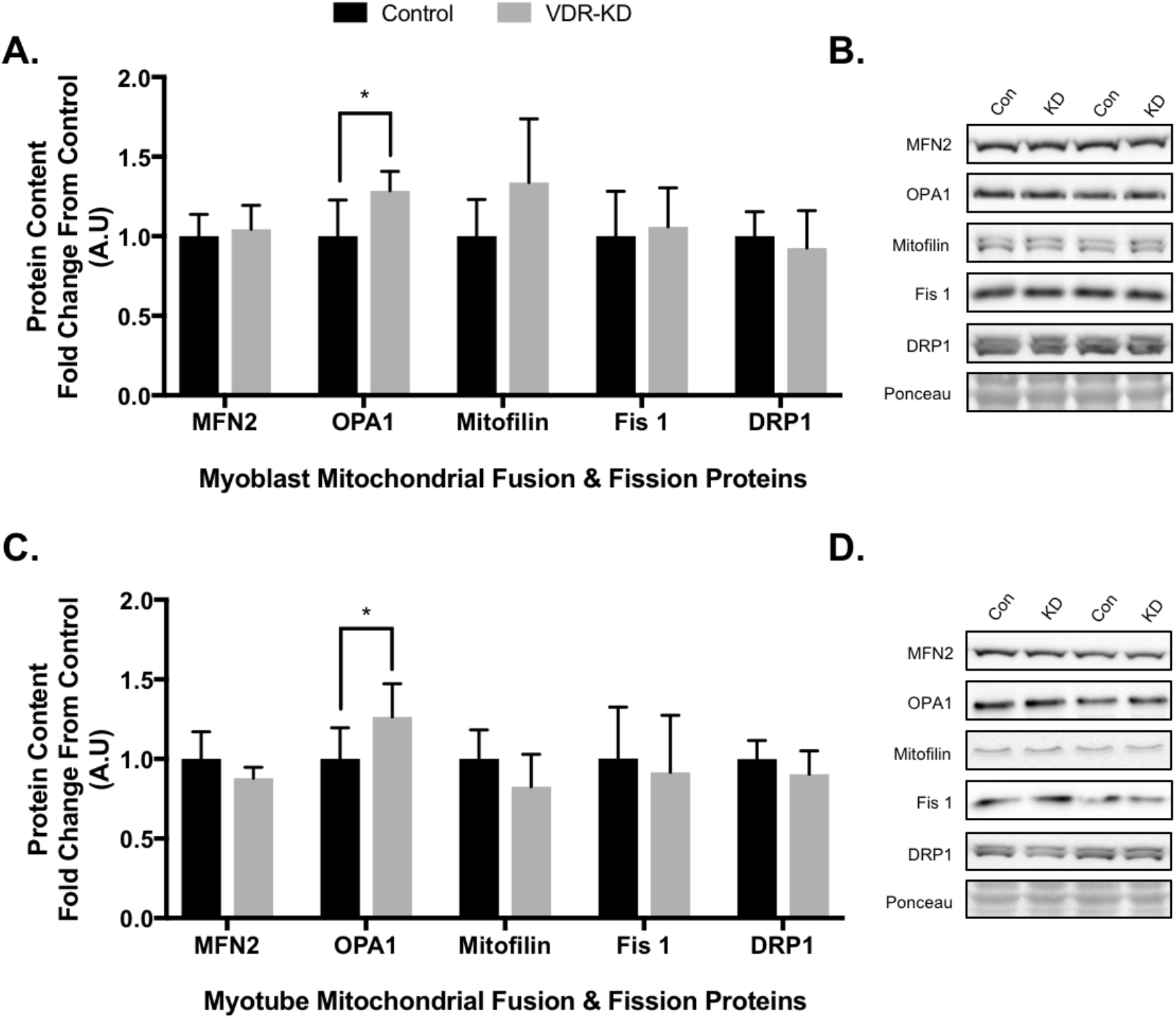
Markers of mitochondrial fission remain unchanged whilst OPA1 protein abundance is increased in VDR-KD myoblasts compared to control. A: Protein abundance of markers of mitochondrial fusion (MFN2 and OPA1) and fission (Mitofilin, Fis1 and DRP1). ^*^*P* < 0.05, independent t-tests. Data mean ± SD (n=6/group) and represented as a fold change from control.

## DISCUSSION

The role of vitamin D within skeletal muscle has received considerable interest in recent years, with current evidence suggesting that vitamin D related metabolites promote mitochondrial function within skeletal muscle (20–24). Building upon previous studies, we demonstrate that loss of VDR function results in significant reductions in mitochondrial respiration in both myoblasts and myotubes (Fig. 2A-D). Furthermore, we report that impairments were specifically observed in respiration derived from oxidative phosphorylation (ATP_ox_) (Fig. 2E) and were not as a result of decreased mitochondrial related protein content (Fig. 3A-D).

Previously, it has been reported that mitochondrial protein content remains unchanged in both human skeletal muscle myoblasts treated with 1α,25(OH)_2_D_3_ and within the quadriceps of skeletal muscle-specific VDR-KO mice (9, 21). Similarly, we also observed no change in mitochondrial protein content in both VDR-KD myoblasts and myotubes. Despite this, it has been reported that the treatment of both human primary and C2C12 myoblasts with vitamin D metabolites resulted in an increase in mitochondrial function (20, 21, 23). Whilst the observed increases in respiration were abolished following siRNA silencing of the VDR in human primary myoblasts (21), the role of the VDR in basal mitochondrial regulation is unknown. Therefore, our results build upon previous findings and indicate that the VDR is required for the maintenance of optimal mitochondrial respiration in myoblasts and myotubes. Furthermore, our results demonstrating that VDR-KD cells have significant reductions in ATP_ox_, suggesting that impairments are intrinsic to the mitochondria following VDR loss-of-function and are not mediated by decreases in mitochondrial protein content *per se*. Despite *in vitro* evidence indicating vitamin D and the VDR regulate skeletal muscle mitochondrial function (20, 21, 23), *in vivo* evidence is currently lacking. Whilst the supplementation of vitamin D has been shown to improve symptoms of fatigue and indirect measures of mitochondrial function (24), further examination of the role of the VDR *in vivo* is warranted.

Given that mitochondria exist within a reticulated network in skeletal muscle (26), we also examined multiple markers of mitochondrial dynamics to ascertain whether loss of VDR function may alter mitochondrial morphology. Whilst we observed no differences in the abundance of MFN2, we did observe small but significant (~15%) increase in OPA1 protein abundance in both VDR-KD myoblasts and myotubes. OPA1 is known to modulate fusion of the inner mitochondrial membrane, cristae remodelling and reduce mitochondrial fragmentation in protection from apoptosis (4, 5, 7). Given the observed impairments in mitochondrial function and membrane potential following VDR-KD, an increase in OPA1 may be a compensatory mechanism to try and rescue mitochondrial dysfunction. Interestingly, OPA1 was also shown to be responsive to 1α,25(OH)_2_D_3_ treatment in human skeletal muscle myoblasts suggesting mitochondrial dynamics within skeletal muscle may be influenced by vitamin D status (21). Further examination of the mitochondrial network in VDR-KD via mitochondrial labelling techniques may shed light upon VitD-VDR-OPA1 interactions in this context.

In summary, we report a requirement for the VDR to maintain optimal mitochondrial respiration in C2C12 myoblasts and myotubes. Reductions in mitochondrial function were a result of reduced ATP_ox_ whilst markers of mitochondrial protein content, fusion and fission were unchanged. Whilst vitamin D supplementation has been shown to improve markers of oxidative phosphorylation and symptoms of fatigue in severely vitamin D deficiency humans (24), our results suggest that the VDR plays a fundamental regulatory role on mitochondrial function in skeletal muscle.

## Author Contributions

S.P.A and A.P conceived and designed research; J.J.B and A.A.K generated VDR-KD and control cell lines. S.P.A performed experiments, analysed data, interpreted results and prepared figures. S.P.A, J.J.B, P.J.A and A.P drafted the manuscript. All authors approved the final version of the manuscript.

